# Investigating the deregulation of metabolic tasks via Minimum Network Enrichment Analysis (MiNEA) as applied to nonalcoholic fatty liver disease using mouse and human omics data

**DOI:** 10.1101/402222

**Authors:** Vikash Pandey, Vassily Hatzimanikatis

## Abstract

Nonalcoholic fatty liver disease (NAFLD) is associated with metabolic syndromes spanning a wide spectrum of diseases, from simple steatosis to the more complex nonalcoholic steatohepatitis. To identify the deregulation that occurs in metabolic processes at the molecular level that give rise to these various NAFLD phenotypes, algorithms such as pathway enrichment analysis (PEA) can be used. These analyses require the use of predefined pathway maps, which are composed of reactions describing metabolic processes/subsystems. Unfortunately, the annotation of the metabolic subsystems can differ depending on the pathway database used, making these approaches subject to biases associated with different pathway annotations, and these methods cannot capture the balancing of cofactors and byproducts through the complex nature and interactions of genome-scale metabolic networks (GEMs). Here, we introduce a framework entitled **M**inimum **N**etwork **E**nrichment **A**nalysis (MiNEA) that is applied to GEMs to generate all possible alternative minimal networks (MiNs), which are possible and feasible networks composed of all the reactions pertaining to various metabolic subsystems that can synthesize a target metabolite. We applied MiNEA to investigate deregulated MiNs and to identify key regulators in different NAFLD phenotypes, such as a fatty liver and liver inflammation, in both humans and mice by integrating condition-specific transcriptomics data from liver samples. We identified key deregulations in the synthesis of cholesteryl esters, cholesterol, and hexadecanoate in both humans and mice, and we found that key regulators of the hydrogen peroxide synthesis network were regulated differently in humans and mice. We further identified which MiNs demonstrate the general and specific characteristics of the different NAFLD phenotypes. MiNEA is applicable to any GEM and to any desired target metabolite, making MiNEA flexible enough to study condition-specific metabolism for any given disease or organism.

**Author Summary:** This work aims to introduce a network-based enrichment analysis using metabolic networks and transcriptomics data. Previous pathways/subsystems enrichment methods use predefined gene annotations of metabolic processes and gene annotations can differ based on different resources and can produce bias in pathways definitions. Thus, we introduce a framework, Minimum Network Enrichment Analysis (MiNEA), which first finds all possible minimal-size networks for a given metabolic process/task and then identifies deregulated minimal networks using deregulated genes between two conditions. MiNEA also identifies the deregulation in key reactions that are overlapped across all possible minimal-size networks. We applied MiNEA to identify deregulated metabolic tasks and their synthesis networks in the steatosis and nonalcoholic steatohepatitis (NASH) disease using a metabolic network and transcriptomics data of mouse and human liver samples. We identified key regulators of NASH form the synthesis networks of hydrogen peroxide and ceramide in both humans and mice. We also identified opposite deregulation in NASH for the phosphatidylserine synthesis network between humans and mice. MiNEA finds synthesis networks for a given target metabolite and due to this it is flexible to study deregulation in different phenotypes. MiNEA can be widely applicable for studying context-specific metabolism for any organism because the metabolic networks and context-specific gene expression data are now available for many organisms.

## Introduction

Nonalcoholic fatty liver disease (NAFLD) is the most common cause of liver disease in western countries [1], affecting an estimated 25% to 45% of the general US population, though this percentage is much higher among individuals suffering from obesity and diabetes [1, 2]. There are two categories of NAFLD: steatosis, which is an accumulation of fat in the liver, and nonalcoholic steatohepatitis (NASH), which consists of the additional presence of liver inflammation and hepatocellular injury with or without fibrosis [2].

Gene deregulation occurs when the cell no longer maintains precise control over certain genes, and therefore expressed proteins, and is associated with various diseases such as NAFLD. Determining which genes are deregulated in disease and how these genes are deregulated can increase the understanding of any disease as well as provide new pathways for therapeutic treatments. To study gene deregulation, pathway enrichment analysis (PEA) uses maps of metabolic processes and subsystem reactions to determine if a given list of genes (gene set) is associated with a certain biochemical pathway, shedding important conceptual insight into gene deregulation. Many PEA algorithms, such as ConsensusPathDB [3] and Piano [3, 4], have been developed to identify biological processes based on gene sets, while other algorithms, such as IMPaLA [5], are based on both gene and metabolite sets. While these algorithms can successfully provide insights about various disease phenotypes, they use predefined metabolic maps that are limited to our current knowledge of these hugely intricate pathways and can differ depending on the pathway database used, meaning they could miss information about the wide range of complex metabolic interactions. These approaches are also subject to biases associated with the different pathway annotations, and they cannot capture the balancing of cofactors and byproducts through the complex nature and interactions of genome-scale metabolic networks (GEMs).

To overcome this, a method has been developed based on elementary flux modes (EFMs), which are non-decomposable flux distributions in metabolic networks [6]. These flux distributions indicate a conceptual description of metabolic pathways. This method computes EFMs from a given GEM and uses tissue-specific transcriptomics to identify a subset of tissue-specific EFMs. However, the enumeration of all EFMs in a GEM can be computationally intractable, and EFMs are not necessarily specific to a target metabolite synthesis, meaning that targeted pathways are not always able to elucidated using EFMs.

One could also use graph-based methods (GBM) that require network topologies and gene expression profiles as inputs to extract deregulated subnetworks that occur between two conditions [7-9]. These methods have been applied to protein-protein networks for prostate cancer study, and they are potentially applicable to metabolic networks. GBM methods use graph-theoretic properties on network topologies but miss information about additional constraints, such as mass balance [10] and thermodynamics [11]. Ideally, the results for these studies would include a set of mass balanced subnetworks that could be used to understand the carbon, energy, and redox flows from precursor metabolites to target metabolites and complex metabolic tasks.

Here, we propose a method called **Mi**nimal **N**etwork **E**nrichment **A**nalysis (MiNEA) that compares two conditions using transcriptomics, proteomics, and metabolomics to identify deregulated minimal networks (MiNs), which are reactions pertaining to various metabolic subsystems that can synthesize a target metabolite. The MiNEA algorithm works by formulating metabolic tasks (MTs) to mimic the various NAFLD phenotypes, such as lipid droplet formation, lipoapoptosis, liver inflammation, and oxidative stress. These MTs include the known set of metabolites necessary to form the modeled phenotype. The cell can use different pathways to form the same metabolites and create the same phenotypes, and the purpose of MiNEA is to identify deregulated alternative routes for a given MT between two conditions, such as a control and treatment. First, MiNEA uses these MTs to compute alternative thermodynamically feasible MiNs applying thermodynamic constraints [11, 12] to a mouse GEM [13]. Then, it uses mouse and human liver sample expression data [1, 14] and identifies deregulated metabolic processes in mice and humans that potentially lead to the NAFLD phenotypes.

MiNEA identified an upregulation in the oxidative stress synthesis network in mice, but this network was found to be downregulated in humans. The cholesterol and triacylglycerol synthesis networks were deregulated in humans only, while the ceramide synthesis network was only deregulated in mice. We found downregulated reactions in the synthesis network for cholesteryl esters, cholesterol, and alanine in both humans and mice. We further identified downregulated reactions in the superoxide anion (SOA) synthesis network in humans, specifically in NASH as opposed to steatosis, while upregulation was found in the SOA synthesis network in mice. This perturbation through the SOA synthesis network in NASH suggested an unbalanced ceramide synthesis, and studies have shown that the ceramide is a key regulator of apoptosis and promotes fibrosis in the hepatic steatosis model [15, 16].

MiNEA can generate MiNs for any target metabolite, e.g. a metabolite produced under specific phenotypes or a biomass building block [17] that is needed for cell growth. Additionally, MiNEA can integrate condition- and context-specific omics data to understand the deregulated phenotypes associated with a set of differentially expressed genes. These characteristics make MiNEA a versatile tool for exploring and understanding different metabolic phenotypes.

## Results and Discussions

### Experimental details for MiNEA

Nonalcoholic fatty liver disease (NAFLD) has been defined as a metabolic disease associated with insulin-resistance syndrome [18]. To study the differences seen in NAFLD phenotypes between mouse and human manifestations, we collected human expression data from the three diagnosis groups, normal (N), steatosis (S), and nonalcoholic steatohepatitis (NS), and mouse expression data from control and DDC-supplemented diet conditions for three genetically different mouse strains, AJ, B6, and PWD. DDC-supplemented diet reproduces steatosis and NASH phenotypes (see Materials and Methods for detail). We refer to these data as *human expression data* and *mouse expression data* throughout this section and integrate it into a mouse model iMM1415 that is constructed based on a human GEM Recon1 [13]. We analyzed both human and mouse MTs using iMM1415 with the assumption that both types of mammalian cells have a similar metabolism (See S1 text). In the following sections, we analyzed the deregulation of minimal networks in core reactions and enriched minimal networks based on the deregulation (up-or downregulated) genes.

### Minimal networks

GEMs represent an entire cellular metabolic network in the form of mathematical constraints [19]. These network reconstructions have grown rapidly in the last decades, and now many GEMs for different organisms are available [20]. GEMs and condition-specific experimental data can be employed to generate and test hypotheses using the Minimal Network Enrichment analysis (MiNEA) framework that has been developed in this study. As compared to a method the had been proposed for finding subnetworks from a GEM that optimizes only some part of metabolic networks to find subnetworks [21], MiNEA optimizes whole metabolic networks to find minimal networks.

To examine the data that can be derived from MiNEA in terms of NAFLD, we first wanted to compute MiNs for each of the MTs in Table 7 (see Materials and Methods for the table) that were significant phenotypes in NAFLD. Instead of representing a main linear route between precursors and targets of MTs, a MiN is instead composed of many subsystems integrated into subnetworks that must be active, meaning reactions of subnetworks carries fluxes, to fulfill the target MT. The MiNEA algorithm facilitates enumeration of alternative MiNs and provides more flexibility to the analysis of different metabolic phenotypes and their respective environmental and genetic perturbations. The use of MT for analysis with MiNEA allows this method to be more easily generalized and applied to the study of other metabolic phenotypes and diseases.

A summary of the MiNs calculated for the MTs in Table 7 (see Materials and Methods for the table) are shown in Table 1. The shortest MiNs were for the synthesis of hydrogen peroxide (H_2_O_2_) (network size = 37), and the longest were for the synthesis of cholesteryl ester (network size = 131; Table 1). We found the greatest number of possible alternatives for the hexadecanoate (HDCA) synthesis and the least number possible for the phosphatidylserine (PS) synthesis (Table 1), suggesting that the HDCA synthesis is more flexible and the PS synthesis less flexible towards alternative formation compared to the rest of the MTs. Reactions that overlapped between all alternative MiNs were called high-frequency reactions (HFRs), and the percentage of HFRs shows how similar or divergent a MiN is compared to the other MiNs. The percentage of HFRs was between 62% to 91% across the MTs (Table 1). In the superoxide anion (SOA) and the hydrogen peroxide (H_2_O_2_) synthesis, 62% and 89% of the reactions were HFRs, respectively, which suggests that the alternative MiNs for a SOA synthesis were more divergent compared to a H_2_O_2_ synthesis.

**Table 1.** Summary of MiNs found for the target metabolites comprising the various NAFLD phenotypes. Synthesis of a target metabolite represents a MT. For each MT, we enumerated alternative MiNs. The number of reactions in each MiN represents the size of the MiN. “# Alt” and “# MT HFRs” represent the number of alternatives and the number of overlapping reactions between all alternatives of a given MT, respectively. The symbol “#Phenotypic HFRs” represents the number of overlapping reactions form all alternatives of a set of MTs that are associated with a given phenotype.

| Phenotypes | Metabolic Tasks (MTs) | Short Name | Min Size # rxns | # Alt | # MT HFRs | % MT HFRs | # Phenotypic HFRs |
| --- | --- | --- | --- | --- | --- | --- | --- |
| Oxidative Stress | Hydrogen peroxide | H<sub>2</sub>O<sub>2</sub> | 37 | 126 | 33 | 89 | 16 |
|  | Superoxide anion | SOA | 42 | 144 | 26 | 62 |  |
| Apoptosis and Inflammation | Alanine | ALA | 98 | 39 | 84 | 86 | 36 |
|  | Ceramide | CRM | 115 | 8 | 105 | 91 |  |
|  | Glutamine | GLU | 116 | 34 | 83 | 72 |  |
| Lipid droplet | Cholesterol | CHOL | 121 | 60 | 108 | 89 | 27 |
|  | Cholesteryl ester | CHOL_ES | 131 | 237 | 112 | 85 |  |
|  | Diacylglycerol | DAG | 82 | 232 | 67 | 82 |  |
|  | Hexadecanoate | HDCA | 75 | 266 | 61 | 81 |  |
|  | Phosphatidic acid | PA | 83 | 245 | 65 | 78 |  |
|  | Phosphatidylinositol | PAIL | 94 | 237 | 68 | 72 |  |
|  | Phosphatidylethanolamine | PE | 128 | 7 | 103 | 80 |  |
|  | Phosphatidylserine | PS | 125 | 6 | 98 | 78 |  |
|  | Triacylglycerol | TAG | 84 | 257 | 62 | 74 |  |

As an expanded example, a MiN from the H_2_O_2_ synthesis comprises many reactions from various metabolic subsystems (Fig. 1). Each MiN represents a group of active reactions, which are reactions that carries fluxes, within multiple metabolic subsystems/pathways that are required for a MT, while a pathway is a group of annotated reactions. In the example in Figure 1, most of the active reactions for the H_2_O_2_ synthesis are from the pentose phosphate pathway (PPP) and the glycolysis/gluconeogenesis pathway. PEA differs in that it identifies a marked deregulation in a pathway as compared to MiNEA, which identifies deregulation in a MiN that could include multiple pathways.

**Figure 1.**
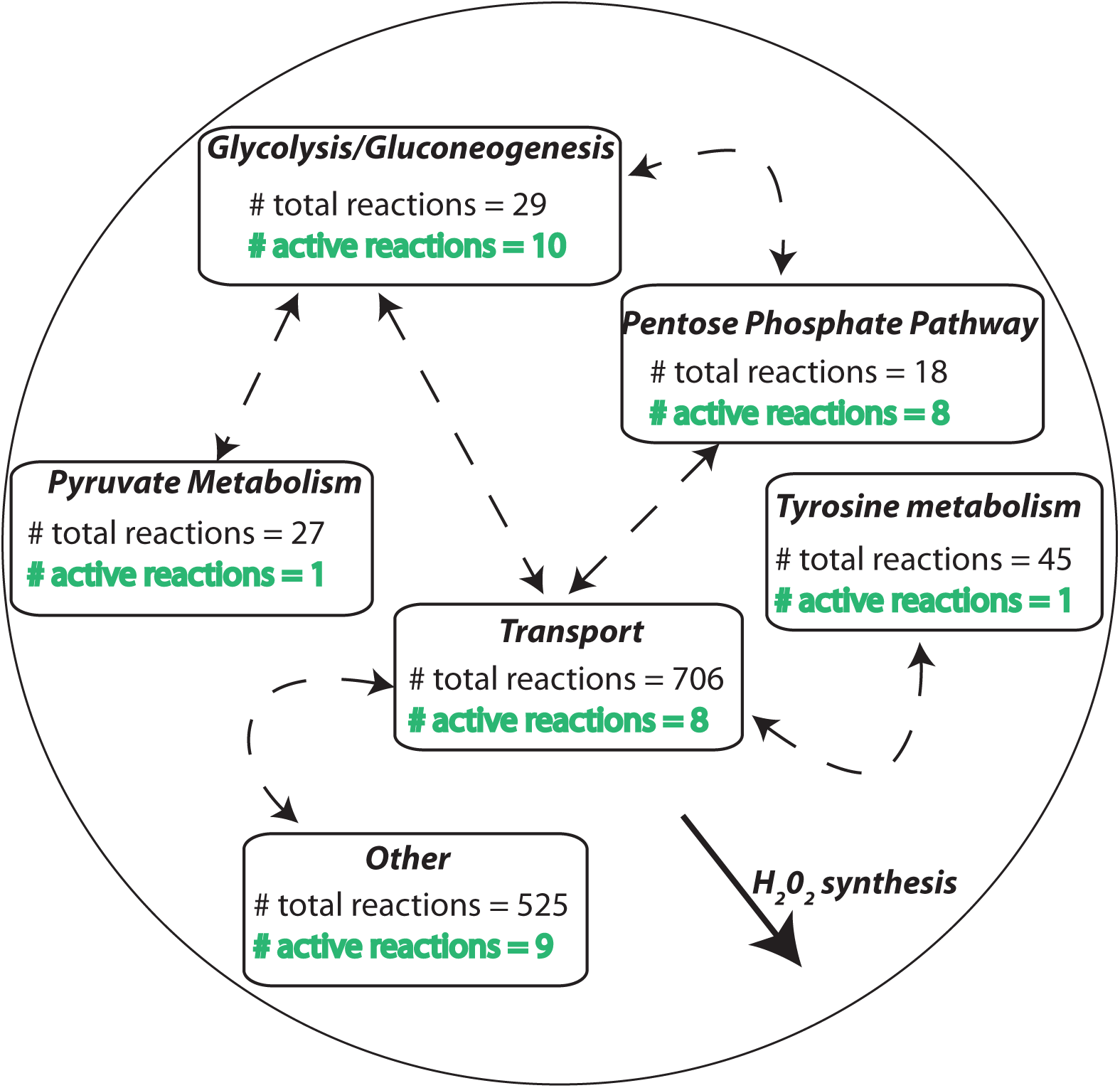
A minimal network (MiN) and high-frequency reactions (HFRs). A MiN of H_2_O_2_ synthesis is illustrated. Each box represents a different metabolic subsystem/pathway. The number of total reactions listed in each box represents the annotated reactions for that subsystem. Active reactions carry non-zero flux, and the number of these present in each subsystem is shown in green text.

We further analyzed the HFRs within each phenotype and found that the most represented metabolic pathway in the apoptosis and inflammation (AI) phenotype was related to amino acid metabolism (# phenotypic HFRs=36; Table 1; Fig. 2).

**Figure 2.**
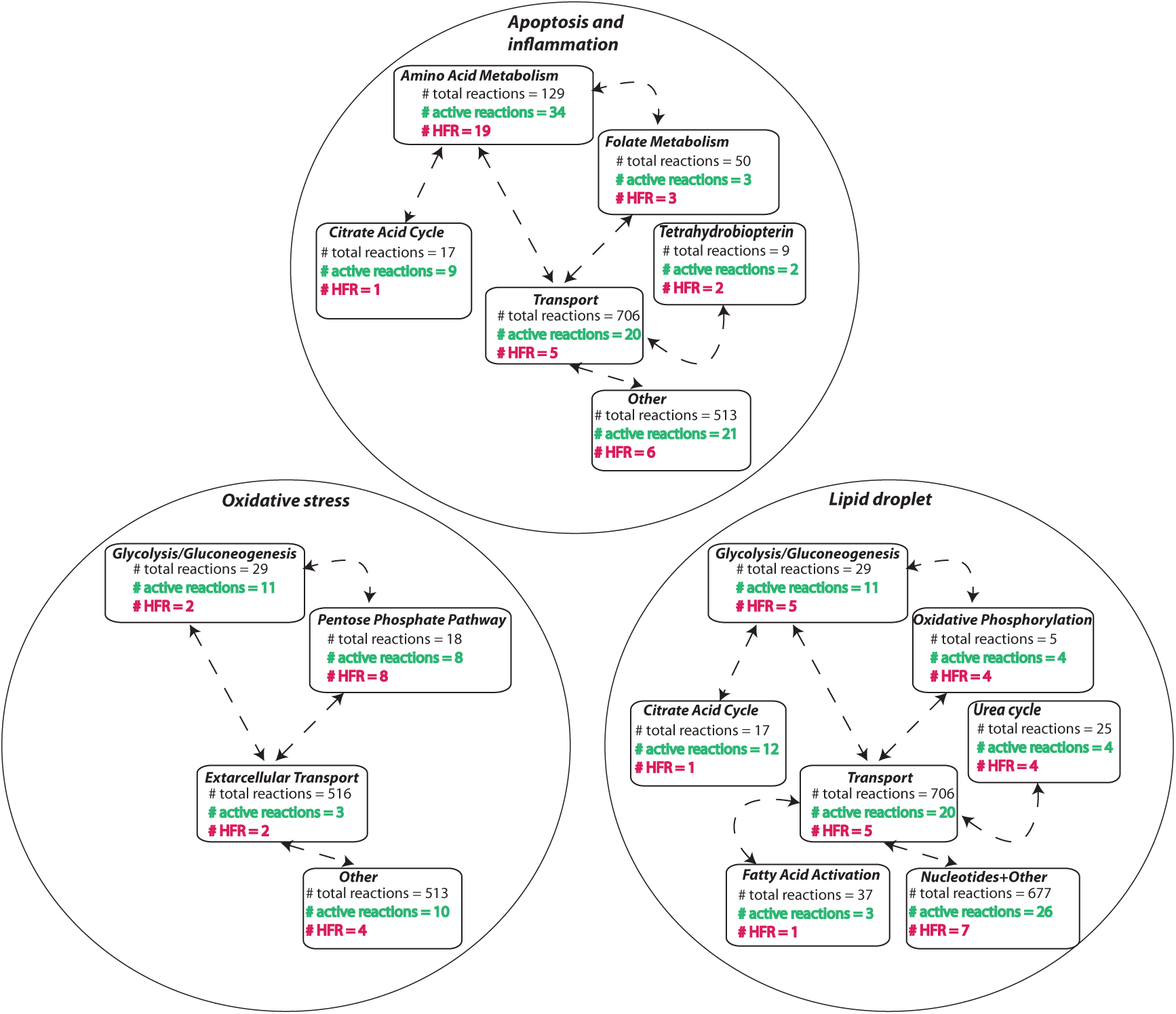
HFRs for all the phenotypes. The number of HFRs in the various pathways is shown for the oxidative stress, apoptosis and inflammation, and lipid droplets phenotypes. See Figure 1 caption for a detailed explanation of the chart.

Interestingly, while the lipid droplet (LD) phenotype has many associated MTs, it shares 27 reactions in which 5 reactions are from the glycolysis/gluconeogenesis pathway (Fig. 2). These shared reactions can be constitutive candidates for the LD phenotypes. The HFRs forming the subsystems in Figure 2 are potential candidates for the key regulators of the NAFLD phenotypes.

### Deregulation in HFRs

The HFRs within each MT create a topology of reaction hubs in metabolic networks that can be similar to protein hubs in protein-protein interaction networks [22, 23], and the misregulation of these hubs in disease can explain the origin of disease phenotypes. Therefore, we performed an analysis of HFR deregulation to study the NAFLD-specific phenotypes. We identified the up- and downregulated reactions from mouse and human expression data (described in S1) and analyzed the HFRs for which the associated genes were found to be deregulated. In humans, the number of downregulated HFRs was higher compared to upregulated HFRs for the NS *vs* N and NS *vs* S across all phenotypes (Table 2). In mice, however, the NS *vs* N, NS *vs* S, and S *vs* N comparisons showed a higher number of upregulated HFRs compared to downregulated ones for the oxidative stress (OS) phenotype, but for the LD and AI phenotypes, we observed similar patterns of up- and downregulated HFRs to humans (Table 2).

**Table 2.**
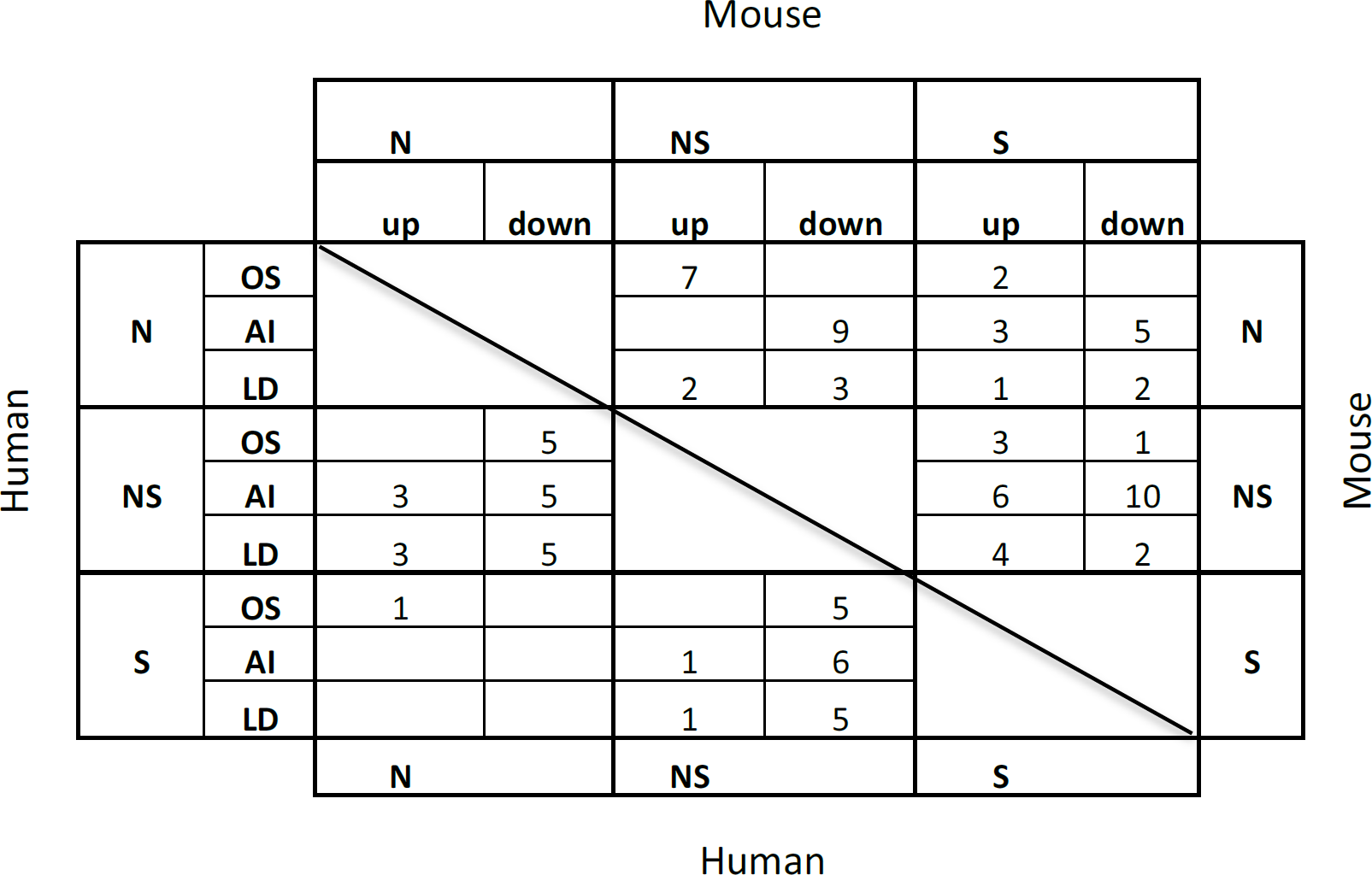
The deregulated HFRs across all comparisons. Numbers in the table indicate the number of up- and downregulated HFRs. In mice, N represents the control diet (N≈AJ, N≈PWD, or N≈B6 strain fed the control diet). Under the DDC-supplemented diet, AJ mice tended towards NASH phenotypes and PWD mice tended towards steatosis phenotypes. Thus, in mice, the symbols NS stand for AJ and S stand for the PWD, all with DDC-supplemented diet (NS≈AJ with DDC-supplemented diet; S≈PWD DDC-supplemented diet).

We further analyzed the deregulation in reaction hubs for each MT to identify the most perturbed MTs in NS and S and the differences between NS and S. In humans, the percentage of downregulated HFRs was higher than upregulated ones for the synthesis of all metabolites in the NS *vs* N and NS *vs* S states, while the percentage of upregulated HFRs was higher than downregulated ones for the S *vs* N state (Fig. 3).

**Figure 3.**
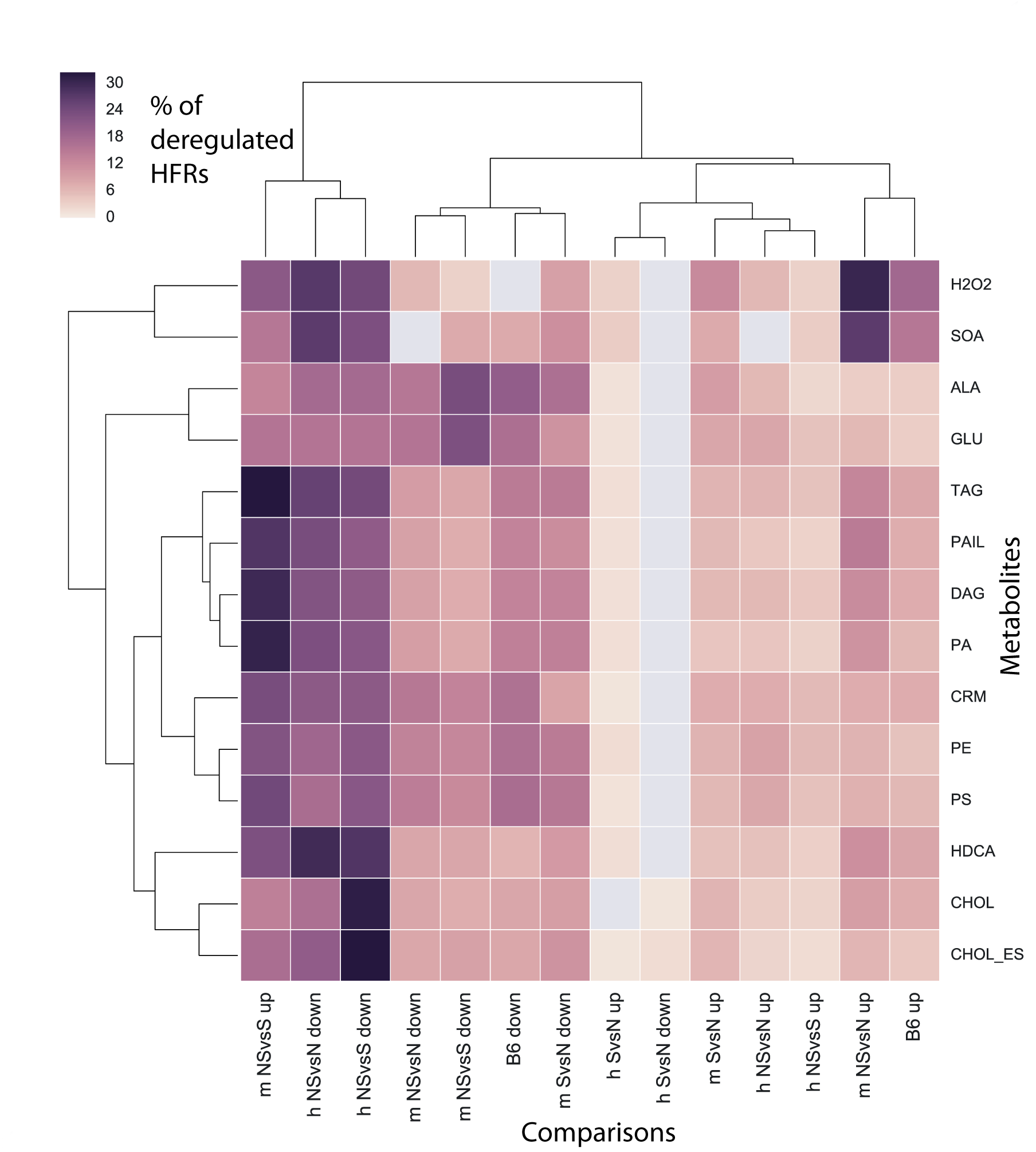
Quantification of up- and downregulated HFRs. Symbols ‘h’ and ‘m’ in comparison’s labels represent human and mouse, respectively, and ‘up’ and ‘down’ indicate up- and downregulation, respectively. See the legend of the Table 3 for the association of mouse strains to the symbols N, NS, and S. “B6 up” and “B6 down” represent the up- and downregulated HFRs for the B6 mouse strain comparing the DDC *vs* control diet. We quantified the percentage of up- and downregulated HFR using a colored heat map.

There was a higher percentage of downregulated HFRs for the cholesterol (CHOL) and cholesteryl ester (CHOL_ES) synthesis networks in NS *vs* S and for the superoxide anion (SOA) and H_2_O_2_ synthesis networks in NS *vs* S (Fig. 3). This means that downregulated HFRs have a higher impact on NASH phenotypes.

Since the mouse strains AJ and PWD show phenotype specific to NS and S, respectively, and the mouse strain B6 show low S and NS phenotypes after feeding with the DDC diet (Materials and Methods) and thus we analyzed B6 separately. For mice, in NS *vs* N (the comparison associated to AJ) and B6 DDC *vs* N comparisons, we found that the percentage of downregulated HFRs in the SOA and H_2_O_2_ synthesis networks was higher than for upregulated ones (Fig. 3). This indicates synthesis networks of oxidative stress were found to be perturbed for AJ and B6 mice.

Additionally, for the NS *vs* S in mice, an elevated percentage of upregulated HFRs was identified for the synthesis of the triacylglycerol (TAG), phosphatidylinositol (PAIL), diacylglycerol (DAG), and phosphatidic acid (PA) metabolites, and a high percentage of downregulated HFRs was found for the alanine (ALA) and glutamine (GLU) synthesis networks (Fig. 3). This is consistent with the observation of lipid droplet perturbation in NASH and steatosis [24], and it further suggests that the synthesis of TAG, PAIL, DAG, and PA are important for lipid droplet formation.

For the synthesis of ROS, such as H_2_O_2_, the human NS *vs* N case presented with a high percentage of downregulated HFRs, while the same case in mice presented with a high percentage of upregulated HFRs (Fig. 3). The reactions of the PPP associated with genes TKT1, TKT2, GND, and TALA were identified as downregulated HFRs in the NS *vs* N human. However, for the NS *vs* N mouse and B6 DDC *vs* N these reactions were identified as upregulated. PPP is known as a source of nicotinamide adenine dinucleotide phosphate (NADPH) that prevents oxidative stress. Thus, the different patterns of regulation observed for these reactions can unbalance both NADPH concentration and H_2_O_2_ synthesis. Interestingly it has been reported that the unbalance in NADPH production through PPP by either an under or over-production of NADPH may induce oxidative stress [25, 26].

The acetyl CoA C acetyltransferase (ACACT1rm) and phosphoglycerate mutase reactions of the ceramide synthesis networks were identified as downregulated HFRs for the NS *vs* S and NS *vs* N in humans along with the NS *vs* N in mice. Thus, perturbation in the genes associated with these reactions can affect the ceramide level in NASH in both mice and humans. In line with this finding, a study suggested that deregulated ceramide production promotes liver injury and the development of NASH through disruption of endoplasmic calcium homeostasis, as well as through the inhibition of autophagy [27].

### Enriched MTs in human and mouse based on deregulated genes

We want to identify deregulation at network level based on an enrichment analysis with deregulated enzymes or genes. A Minimal network (MiN) of a metabolic task (MT) contains metabolites, active reactions and their associated enzymes in order to fulfill the MT. To identify deregulated MTs and their associated MiNs (see Materials and Methods section for definitions) in steatosis and NASH, we performed Minimal Network Enrichment Analysis (MiNEA) (see Materials and Methods section) using mouse and human expression data. We identified deregulated genes (S1 text; Table S1-S3), and for each set of deregulated genes, we estimated the *p* value.

Applying MiNEA to the NS *vs* N human case, we identified deregulated MiNs which corresponded to the synthesis of H_2_O_2_, PA, and TAG (Table 3; Table S4 & S5). We computed the percentage of significantly enriched MiNs from all generated alternatives of each MT and scored the deregulated MiNs using the **A**lternative **Mi**nimal **N**etwork **F**requency (AMiNF), which represents the percent of MiNs with enrichment in deregulated genes. For the NS *vs* N case in humans, H_2_O_2_ synthesis had the highest AMiNF (AMiNF = 0.143) compared to all other deregulated MTs (Table 3), suggesting that the deregulation of H_2_O_2_ metabolism and oxidative stress contributes to NASH. Indeed, a study has shown that a marked alteration of H_2_O_2_ concentration can lead to different types of oxidative stress [25].

**Table 3.**
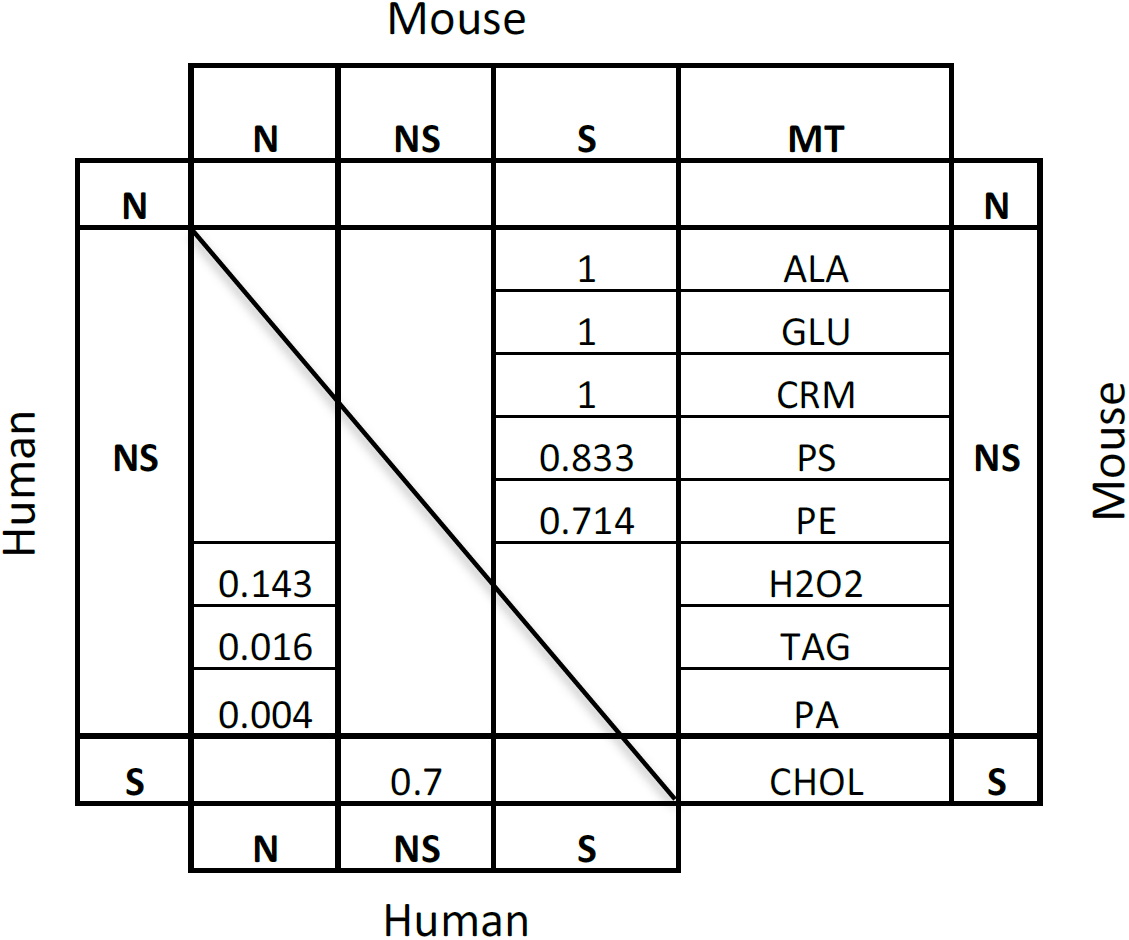
The significantly deregulated MTs across all comparisons based on deregulated genes. The numbers in the table indicate Alternative Minimal Network Frequency (AMiNF). The mouse strains associated with N, NS, and S are described in the legend of Table 2. For the calculation of AMiNF *p* < 0.05 was considered statistically significant.

When we applied MiNEA to NS *vs* S in humans, we found the most deregulated MiNs to be associated with CHOL synthesis (AMiNF = 0.7). Interestingly, TAG synthesis was also deregulated in NS *vs* N, suggesting that lipid droplet formation can be perturbed in the NASH state but through different components.

In mice, we did not observe marked deregulation in NS *vs* N and S *vs* N, but we observed significant deregulation in NS *vs* S for the synthesis of ALA, GLU, ceramide (CRM), PS, and phosphatidylethanolamine (PE) (Table 3), suggesting that these MTs are subject to different perturbations between the NASH and steatosis states.

In humans, MiNEA identified deregulation in the CHOL and TAG synthesis network, which are lipid droplet constituents. Similar deregulation was observed for the lipid droplet constituents in mice through the PS and PE synthesis networks. Thus, because MiNEA was applied with information on deregulated genes, we find that in both humans and mice, lipid droplet formation is perturbed but through different lipid constituents.

### Enriched MTs in humans and mice based on up- and downregulated reactions

To identify whether a specific up- or downregulated MiN was associated with the NAFLD phenotypes, we performed MiNEA while enriching for up- and downregulated reactions separately (S1 Text).

In humans, we identified upregulated MiNs for the synthesis of PS and PE in N *vs* S (Table 4; Table S6 & S7). For the NS *vs* N and NS *vs* S cases, we identified downregulated MiNs associated with the synthesis of CHOL_ES, CHOL, ALA, and GLU (Table 5; Table S8 & S9), but none were found to be upregulated. This implies that in NASH the synthesis of cholesterol and the glucogenic amino acids alanine and glutamine are downregulated. Interestingly, in the NS *vs* N case, only 3% of the cholesterol synthesis alternatives were significantly downregulated (Table 5). Although it is a relatively small frequency, it can provide important leads, such as deregulation in metabolic tasks and hypotheses that would have been missed by the commonly used pathway enrichment methods, such as [3, 5]. In general, these studies suggested that MiNEA can identify deregulated alternative MiNs under different conditions. For example, previous pathway analysis methods [3, 5] lack an enumeration of alternatives that would have been failed to identify cholesterol synthesis as downregulated in NS *vs* N (Table 5).

**Table 4.**
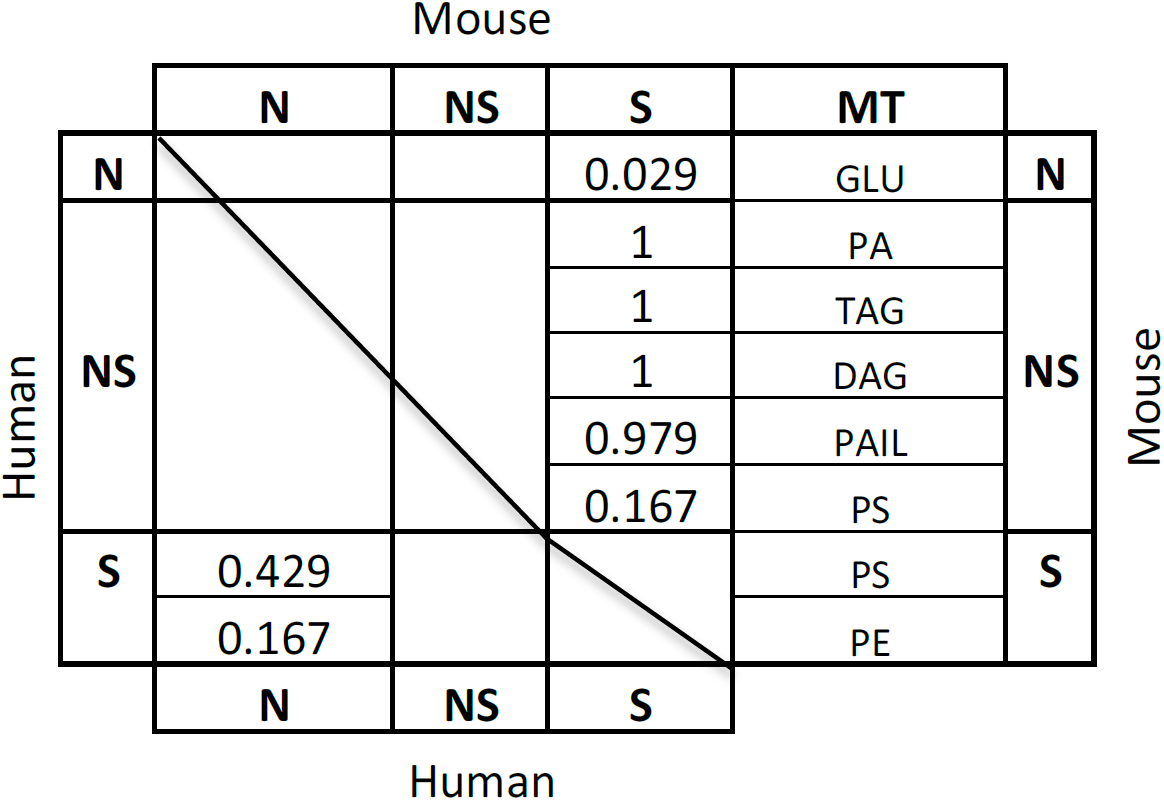
The significantly deregulated MTs across all comparisons based on upregulated reactions. For a detailed understanding, see the legend of Table 2. Numbers of the table represents AMiNF and for the calculation of AMiNF *p* value < 0.005 was used.

In mice, the MiNs associated with the synthesis of ALA, CRM, PE, PS, CHOL_ES, and GLU were identified as downregulated in NS *vs* N (Table 5). Interestingly, in S *vs* N, only the GLU synthesis network was upregulated (Table 4). For the NS *vs* S case, we found major deregulations. Specifically, we found five upregulated networks (PA, TAG, DAG, PAIL, and PS) and five downregulated (ALA, GLU, CRM, PS, and PE) synthesis networks. Within the alternative MiNs that are used for the synthesis of PS, we found some networks that were upregulated and some that were downregulated. Since there was a higher frequency of downregulated MiNs (AMiNF = 0.5) compared to upregulated ones (AMiNF = 0.167), we could hypothesize that the downregulation of PS synthesis is an important molecular mechanism specific to NASH. Here, if we use pathways enrichment methods [3, 5] then we cannot associate two different AMiNF score for a given MT due to the lack of alternative enumeration and would have been failed to give a higher confidence to downregulation of PS synthesis.

**Table 5.**
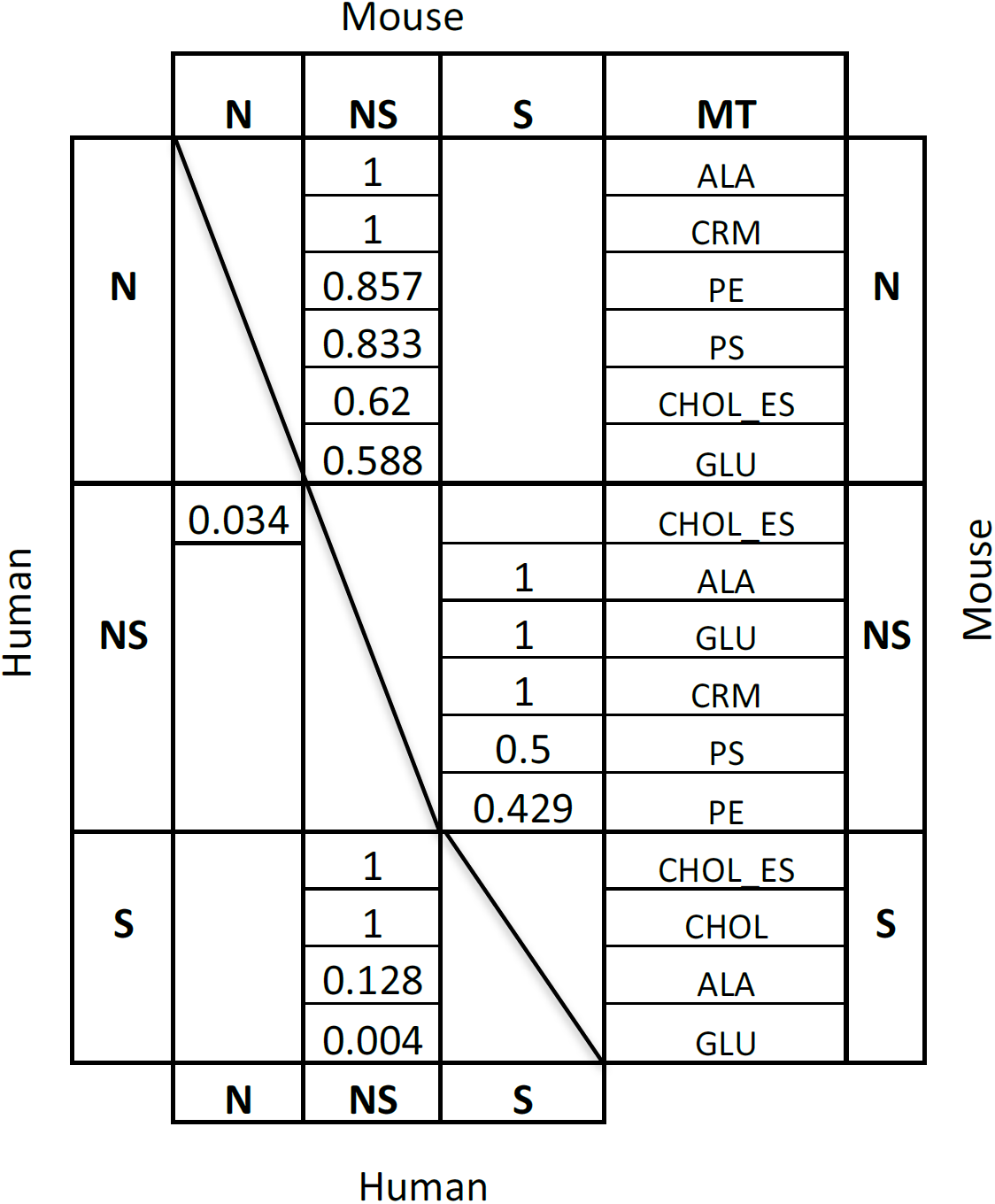
The significantly deregulated MTs across all comparisons based on downregulated reactions. For detail understanding see the legend of Table 4. For the AMiNF calculation *p* < 0.005 was considered statistically significant.

The cholesterol, glutamine, and alanine syntheses were found to be similarly deregulated in humans and mice for the NS *vs* N case. In contrast, the PS synthesis was differently deregulated between the two species (Table 4 and 5). Despite this opposite response in the regulation of the PS synthesis between humans and mice, it appears that the level of PS is perturbed in NASH. Overall this observation suggests that different regulations in gene expression can translate to different effects on the production of lipid droplet synthesis.

### Integration of metabolite concentration data into a thermodynamically feasible metabolic model

The integration of metabolomics can change the size and structure of the MiNs for MTs by eliminating thermodynamically infeasible reaction directionalities. In mice, there are available metabolic profiles for the N, NS, and B6 cases [1]. We integrated these profiles into the iMM1415 [11, 13] and found that, in this case, there are not marked changes in the MiNs and the result in the HFRs, which means we did not found a marked impact on the synthesis networks of NAFLD phenotypes. A further sensitivity analysis can guide future metabolomics studies to target metabolites that have a marked thermodynamic impact on a metabolic network [28]. Thus, the information content of such metabolite sets can be influential and more relevant to the phenotypes under study.

We applied MiNEA for the study of deregulated metabolic processes rather signaling and regulatory processed because metabolism is better characterized, as is shown with the increased availability of GEMS for many organisms. Despite the challenges in reconstructing constraint-based signaling networks, recently such reconstructions have started to become available. Signaling networks for the toll-like receptor (TLR) and epidermal growth factor receptor (EGFR) are now available, and MiNEA can be extended to include these networks [29, 30], meaning that MiNEA could easily be applied to the study of deregulation in signaling networks.

## Materials and Methods

### Microarray gene expression analysis of human liver samples

The microarray gene expression data pertaining to the three diagnostic groups (normal [N], Steatosis [S], and NASH [NS]) were collected from the ArrayExpress public repository for microarray data under the accession number E-MEXP-3291 [14]. We performed an analysis of the differentially expressed genes (DEGS) and a pairwise comparison between diagnostic groups. To control the false discovery rate at level of 0.05, multiple hypothesis testing was used [31].

### Mice phenotypes after feeding a 3,5-diethoxycarbonyl-1,4-dihydrocollidine (DDC)-supplemented diet

NASH phenotypes can be reproduced in mouse models by treatment of chronic intoxication of with 3,5-diethoxycarbonyl-1,4-dihydrocollidine (DDC)-supplemented diet [1]. Three genetically different mouse strains AJ, B6, and PWD were fed with a DDC-supplemented diet. Under DDC-supplemented diet, the steatosis and NASH phenotypes were the most obvious in the PWD and AJ strains, respectively, while for the B6 strain, low levels of the steatosis and NASH phenotypes were observed [1]. The B6 mouse strain showed less induction of the NASH phenotype than the AJ strain [1].

### Association of mouse strains with human diagnoses groups

The normal (N) feeding group is associated with mice fed the control diet. All mice strains, AJ, B6 and PWD, were fed this diet and are labeled as N≈AJ, N≈PWD or N≈B6 to indicate the control diet and mouse strain. After feeding the DDC-supplemented diet, AJ and PWD showed high NASH phenotypes and high steatosis phenotypes, respectively [1]. Thus, under the DDC-supplemented diet, the symbol for steatosis (S) is associated with the PWD mouse strain and the symbol for NASH (NS) is associated the AJ mouse strain (with DDC-supplemented diet: NS≈AJ, S≈PWD). Thus, NS *vs* N represents the AJ mouse strain treated with the DDC diet *vs* the AJ mouse strain fed the control diet, S *vs* N represents the PWD mouse strain treated with the DDC diet *vs* the PWD mouse strain fed the control diet, and NS *vs* S represents the AJ mouse strain treated with the DDC diet *vs* the PWD mouse strain fed the DDC diet.

### Metabolite concentration and RNA-seq gene expression data from mouse liver samples

Metabolite concentration data and RNA-seq gene expression data from mouse liver samples of the three mouse strains (AJ, B6, and PWD) measured under the control and DDC-supplemented diet conditions were collected from the work of Pandey et al. [1]. The R-package “edgeR” [32] was used to identify the differentially expressed genes (DEGs) using three biological replicates of each mouse strain for both the control and the DDC-treated conditions. Strain-wise identification of the DEGs between the DDC-treated and control states was performed separately for each mouse strain. We used the Benjamini-Hochberg procedure implemented in edgeR to control the false discovery rate to a level of 0.05 [31].

### Formulation of metabolic tasks (MTs) based on steatosis and steatohepatitis (NASH) phenotypes

Steatosis can occur due to the accumulation of lipid droplets in the liver, which can subsequently lead to NASH upon liver inflammation [33]. Lipid droplets are composed of many lipid metabolites [34, 35], which are summarized in Table 7. Reactive oxygen species (ROS), such as the superoxide anion, can also damage hepatic membranes and play an important role in the development of NASH [36]. To study these various phenotypes and their role in the various forms of liver disease, the metabolites associated with their various phenotypes were examined. Table 7 summarizes the key metabolites associated with the major liver disease phenotypes: lipid droplets, liver inflammation, apoptosis, and oxidative stress. We investigate the synthesis networks of these metabolites using MiNEA, where we called the synthesis of a metabolite as a metabolic task (MT).

**Table 7:** Metabolites associated with NAFLD phenotypes. This provides a summary of key metabolites associated with the four phenotypes and their corresponding references. Metabolites are taken from the mouse metabolic model of Sigurdsson and colleagues [13].

| Metabolites | Phenotype | Reference |
| --- | --- | --- |
| **Diacylglycerol, phosphatidylethanolamine, cholesteryl esters, triacylglycerol, hexadecanoate, cholesterol, phosphatidic acid, phosphatidylinositol, phosphatidylserine** | Lipid droplets | [34, 35, 37] |
| **Ceramide** | Inflammation | [27, 37, 38] |
| **Alanine, glutamine, ceramide** | Apoptosis | [39] |
| **Superoxide anion, hydrogen peroxide** | Oxidative stress | [36] |

### Minimal network enrichment analysis (MiNEA) algorithm

The cell can use various different pathways depending on the current state and conditions of the environment to fulfill its immediate metabolic needs. In order for MiNEA to identify deregulated alternative routes for a given MT between two conditions (Fig. 4), the inputs required are a metabolic model, a list of MTs, and gene or protein expression data. In this work, we used a mouse GEM (iMM1415) reconstructed by Sigurdsson and colleagues [13], a list of MTs which are the synthesis of the metabolites shown in Table 7, and gene expression data of mouse and human liver samples [1, 14].

**Figure 4.**
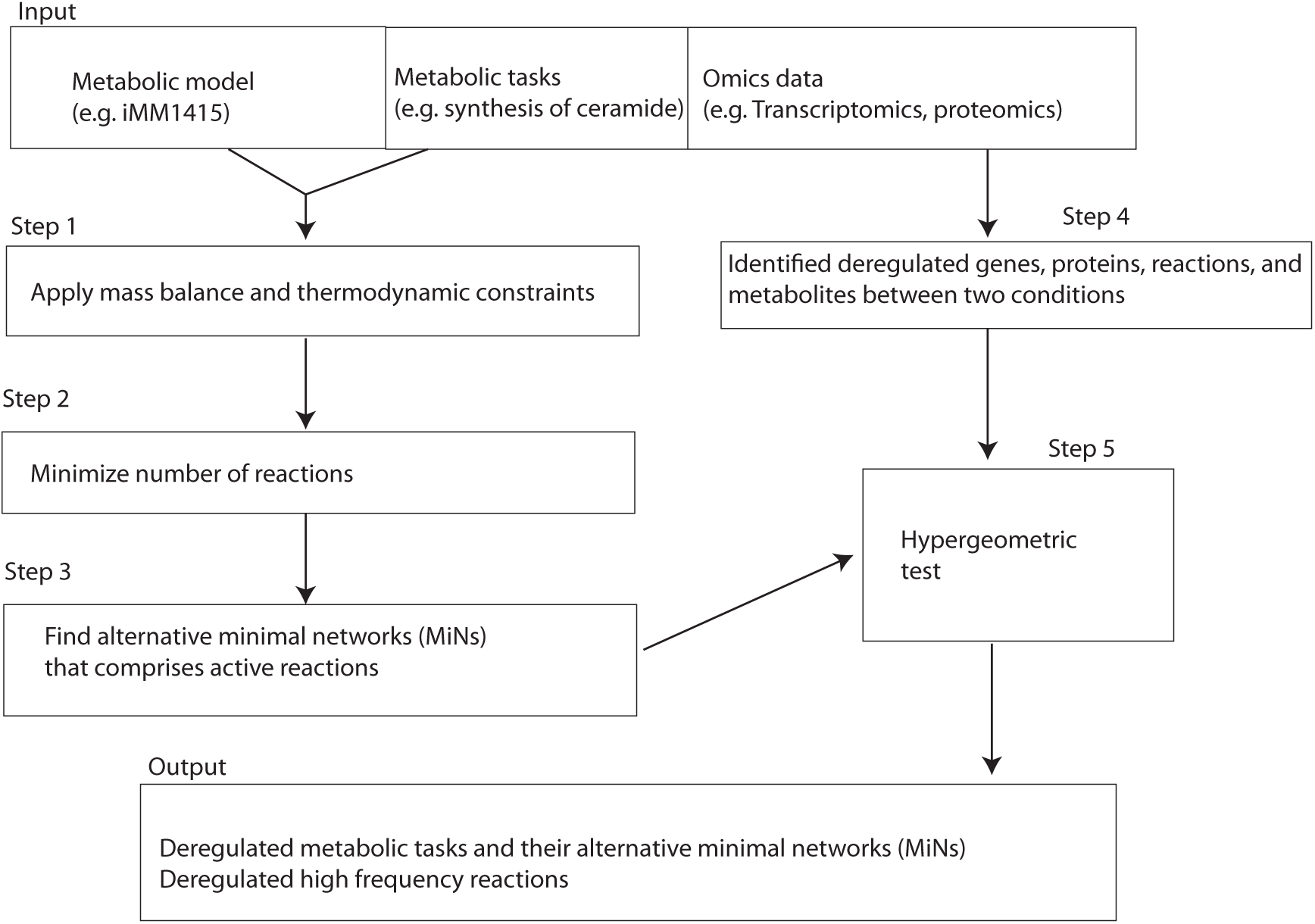
Minimal network enrichment analysis (MiNEA) overview. For each metabolic task (MT), MiNEA computationally enumerates MiNs that comprise active reactions for the MT. The inputs required for the MiNEA analysis are a genome-scale metabolic network (GEM), a given list of metabolic tasks (MTs), and transcriptomics data. Using this, alternative minimal networks (MiNs) are enumerated for MTs using a GEM. (steps 1–3). Transcriptomics data are used to identify differentially regulated genes between two conditions (step 4). To identify deregulated MTs and their associated MiNs, a hypergeometric test is performed with a set of deregulated genes (step 5).

The MiNEA calculations begin by applying thermodynamic constraints to the model as described previously [11] to eliminate thermodynamically infeasible reactions, meaning that reactions from the metabolic network can carry fluxes only if thermodynamics allows [12] (Fig. 4; step 1). Then, it enumerates all the thermodynamically feasible minimal-size networks (MiNs) that are active for the given list of provided MTs (Fig. 4; step 2), and all of these MiNs are composed of reactions that carry non-zero flux. For this step, the following mixed-integer linear programming (MILP) problem is applied with the objective of minimizing the number of reactions that carry flux or maximizing the number of reactions that cannot carry flux, while enforcing that the network should be able to synthesize the metabolites listed in Table 1:
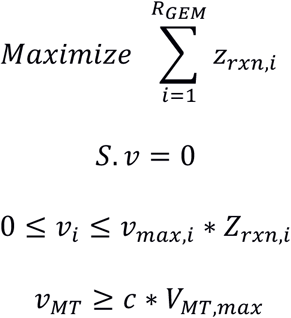
*R*_*GEM*_ is the number of reactions in a GEM and *z*_*rxn,i*_ is a binary variable associated with reaction *i*. Reversible reactions were split into two forward reactions. *V*_*MT,max*_ is the maximum yield to produce a metabolite associated with a MT. The parameter *c* is generates MiNs that allow for flexibility on the yield from the MT. We chose *c* = 0.8 to allow for a yield of at least 80% of the maximum yield for the associated MT.

All alternative MiNs that are the minimum size (msize) for the synthesis of the metabolites in Table 1 are enumerated as described by Figueiredo and colleagues [40] (Fig. 4; step 3). One can enumerate many alternative networks larger than msize, such as msize+1 and msize+2. Each MiN is a subnetwork comprising a set of reactions, metabolites, and reaction-associated genes through gene-protein-reaction (GPR) association. In step 4, the deregulated genes and proteins that differ between the two conditions under study are identified (Fig. 4). Finally, a hypergeometric test is performed on the sets of deregulated genes or deregulated reactions to identify the deregulated MiNs (Fig. 4; Step 5).

### Minimal network significance based on gene set and reaction set (Step 5)

The significance of the MiN based on the deregulated genes was calculated using the hypergeometric probability density function (P),
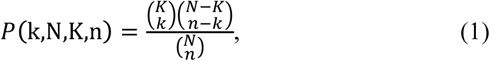
where *k* and K are the numbers of deregulated genes and the number of total genes in a given MiN, respectively, and n and N are the total number of deregulated genes and the total number of genes of a metabolic network, respectively. In this context, the term “deregulated genes” is used for both up- or downregulated genes.

A reaction can have three states: upregulated, downregulated, or unregulated. The regulation of a reaction was determined based on its associated differentially expressed genes. According to this metric, a reaction was identified as upregulated or downregulated if the corresponding genes are only upregulated or only downregulated, respectively. A reaction that is associated with a mixture of up- and downregulated genes is not characterized as regulated due to the inconsistency of gene expression. Up- and downregulated MiNs, which could contain various combination of up- or downregulated reactions, were identified in Step 4 of MiNEA (Fig. 1) based on the total number of up- or downregulated reactions, respectively. As expected, up- and downregulated MiNs comprise markedly high numbers of up- and downregulated reactions, respectively.

The significance of a MiN based on upregulated or downregulated reactions was calculated using multivariate Fisher's hypergeometric distribution. This method has been previously used for the selection of tissue-specific elementary modes using gene expression data [6]. To identify significantly upregulated MiNs in a given set of MiNs, we selected those MiNs that contained an elevated number of upregulated reactions and as few as possible downregulated ones, while for the identification of significantly downregulated MiNs, we selected MiNs with an elevated number of downregulated reactions and as few as possible upregulated ones.

Assume R and T are the total numbers of reactions in a GEM and a MiN, respectively, which can be decomposed as follows:
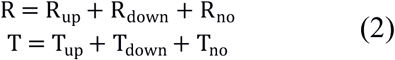
*R*_*up*_, *R*_*down*_, and *R*_*no*_ represent the number of upregulated, downregulated, and unregulated reactions in a GEM, respectively. In the context of a MiN, T is the total number of reactions, and *T*_*up*_, *T*_*down*_, and *T*_*no*_ are the number of upregulated, downregulated, and unregulated reactions, respectively.

To consider a MiN as upregulated, we need to ensure that the pair (*T*_*up*_, *T*_*down*_) in the MiN of *T* reactions cannot arise by chance in the context of the whole network. To obtain a better upregulation by chance, the *p* value was computed using equation (3). Note that an equal or better outcome than the pair (T_up_, T_down_) satisfies two conditions: (i) the number of downregulated reactions is smaller than or equal to T_down_, and (ii) the number of upregulated reactions is greater than or equal to T_up_, whereas the number of reactions in the MiN remains unchanged.
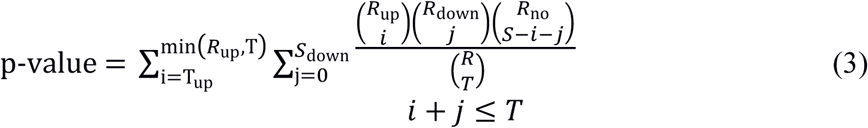

We can compute the *p* value for downregulated reactions the same way as for the upregulated reaction simply by changing ***up*** with ***down*** and ***down*** with **up** in the above equation.

### Deregulated percentage (DRP)

We calculated the deregulated percentage (DRP) for each MiN, which indicates the percentage of up- or downregulated genes in a MiN. For example, if a given MiN comprises 20 genes, and 5 genes of the MiN are deregulated, the then DRP is equal to 0.4. Since the active reactions in a MiN were classified as either up- or downregulated, we calculated both upregulated percentage (UPR) and downregulated percentage (DnRP) for each MiN.

### Alternative minimal network frequency (AMiNF)

To extend the degree of confidence of the results from MiNEA, we introduced the alternative minimal network frequency (AMiNF) metric that determines the significantly deregulated percentage of MiNs for a given MT. If all the alternative MiNs for a MT are significantly deregulated, AMiNF takes the value 1. The AMiNF is 0 when none of the MiNs for a MT are significantly deregulated.

### MiNEA implementation

MiNEA was implemented using Matlab r2016a (The Mathworks, Natick, MA, USA), and MILP problems were solved using CPLEX solver (ILOG, Sunnyvale, CA, USA). The MILP gaps for all problems were converged to less than 0.05% in less than 2,400 s. We also plan to make the MiNEA algorithm available as a tool for distribution to the community.

## Conclusions

In contrast to graph-based methods (GBM) and pathway enrichment analysis (PEA), the methodology introduced here (MiNEA) expands the notion of pathway into in a set of mass balanced subnetworks that can be used to understand the carbon, energy, and redox flows from precursor metabolites to target metabolites and complex metabolic tasks. One of the main advantages of MiNEA compared to PEA is that MiNEA attempts an enumeration of alternative minimal networks for each metabolic task (MT), which helps us understand and study the MT flexibility. Although a large number of alternative enumerations for a complex metabolic network can be time-consuming, once the enumeration is completed, MiNEA applies a statistical analysis that is fast and extracts additional information, such as pertaining to the deregulation of MTs or the deregulation in reaction hubs. As an example of the power of MiNEA, we identified deregulation in key metabolic network of the ceramide and hydrogen peroxide synthesis, of NASH in both humans and mice. We also identified similar deregulation in NASH for the cholesterol synthesis networks in humans and mice, and we found opposite deregulation for the phosphatidylserine synthesis network between humans and mice. MiNEA is highly applicable for the study of context- or condition-specific metabolism because using this one can identify synthesis networks for any given target metabolite and further can employ condition-specific transcriptomics, proteomics, and metabolomics data.

## Supplementary Tables

Table S1. Differentially expressed genes for human and mouse expression data.

Table S2. Upregulated reactions for human and mouse data.

Table S3. Downregulated reactions for human and mouse data.

Table S4. Minimal network enrichment analysis based on differentially expressed genes. Description of the table heading is illustrated in ‘Symbol’ sheet of the excel file. Please see the Symbol sheet for the detail description for the tables: S5-S9.

Table S5. Minimal network enrichment analysis based on upregulated reactions.

Table S6. Minimal network enrichment analysis based on downregulated reactions.

Table S7. Marked deregulated minimal networks selected based on Table S4.

Table S8. Marked deregulated minimal networks selected based on Table S5.

Table S9. Marked deregulated minimal networks selected based on Table S6.

## Acknowledgments

VP and VH are supported by the RTD grant MalarX within SystemsX.ch, the Swiss Initiative for Systems Biology evaluated by the Swiss National Science Foundation. VP and VH are supported by the École polytechnique fédérale de Lausanne.

## Author Contributions

Conceptualization and methodology: VP and VH. Implementation of the method and data analysis: VP. Wrote the paper: VP and VH. Supervision: VH.

## Supporting Information

### Comparison of human and mouse genome-scale metabolic models

Mouse genome-scale metabolic model, iMM1415 [13] was reconstructed based on a human genome-scale metabolic model (Recon 1, [41]). Sigurdsson and collogues found that the mammalian organism with the highest number of genes homologous to Recon 1 genes was the mouse (*Mus musculus*) (1,415 genes, 97%). We compared iMM1415 and Recon1 and found that the iMM1415 shares 98% of reactions with Recon1 and in the reaming 2 % reactions more than 1.5 % reactions were associated to the extracellular transport mechanism. This suggests that Recon1 and iMM1415 have very similar metabolism.

### Identification and analysis of deregulated genes and reactions in human and mouse liver samples

To understand the nonalcoholic fatty liver disease (NAFLD) and the difference of the disease in mouse and human we collected human expression data form the three diagnosis groups: normal (N), steatosis (S), and nonalcoholic steatohepatitis (NS), and mouse expression data form the control and DDC-supplement diet conditions for the three mouse strains: AJ, B6 and PWD (*Materials and Methods*). These data were referred as *human expression data* and *mouse expression data* throughout this section.

### Differential expressed genes in human liver samples

To understand the NAFLD physiology we analyzed the differentially expressed genes (DEGs) in the *human expression data*. Out of the 1415 metabolic genes in iMM1415 [13], we identified 29, 484 and 363 DEGs in S *vs* N, NS *vs* S, and NS *vs* N, respectively (Fig. S1 upper panel and Table S1). Only 29 DEGs between S *vs* N may explain that the metabolic state was very similar between normal and steatosis. The total number of DEGs between NS *vs* N and NS *vs* S were much higher than S *vs* N, and thus suggest a more pronounced alteration of the metabolic state of nonalcoholic steatohepatitis compared to steatosis and normal. Furthermore, the number of downregulated genes is greater than the number of upregulated genes in exclusively NS *vs* N and NS *vs* S (Fig. S1), suggesting that, in human NASH, the perturbation leading to the metabolic state that characterizes it, is reached by downregulated genes.

### Differential expressed genes in mouse liver samples

We analyzed the *mouse expression data* for DEGs as described in the *material and methods* section. Out of the 1415 metabolic genes in iMM1415, the total number of DEGs between control and DDC-supplemented diet was similar across all strains with 247, 248 and 221 for AJ, B6 and PWD, respectively (Fig. S2 lower panel). Here, AJ and PWD are associated with NS *vs* N and S *vs* N, respectively (see materials and methods). Many up- and down-regulated genes were strain-specific. The number of up- and down-regulated strain-specific gene pairs for AJ, B6, and PWD were (48, 20), (33, 29), and (39, 62), respectively (Fig. S1 lower panel and Table S1). The number of up-regulated genes was greater than the number of down-regulated genes for the AJ strain, while for the PWD strain the opposite was true. Numbers of up- and down-regulated genes were very similar for the B6 strain. Interestingly, the observed differences qualitatively correlate with the strains’ phenotypes: steatohepatitis phenotypes were observed high, low, and unspecific for the AJ, B6, and PWD mouse strains, respectively [1].

We identified deregulated genes form the AJ under the DDC-supplemented diet *vs* PWD under DDC-supplemented diet and this is associated with NS *vs* S (see materials and methods). For this comparison we identified 191 and 100 up- and down-regulated genes, respectively (Table S1).

### Up- and downregulated reactions in human and mouse

A reaction is marked as down regulated if genes associated to the reaction is down-regulated and if genes associated to the reaction is upregulated then the reaction is called as upregulated. If a reaction is associated to genes with mix-regulation (up- and down-regulation) then the reaction is not marked with up- or down-regulated. Regulations of reactions are computed for the *human expression data* and *mouse expression data* (Fig. S2).

Only for NS *vs* S, the number of down-regulated reactions was higher than the number of up-regulated reactions (Fig. S2 upper panel), and a similar trend was observed for the number of up- and down-regulated genes (Fig. S1 upper panel and Tables S2-S3). The number of reactions up- and down-regulated was similar for NS *vs* N, but the number of down-regulated genes was higher than the number of upregulated genes (Fig. S1 and S2 upper panels). For S *vs* N, the number of down-regulated reactions was higher than the number of up-regulated reactions; however, the numbers of up- and down-regulated genes for these conditions were similar (Fig. S1 and S2 upper panels). This observation indicates that the reaction regulation (RR) is not always in agreement with the gene regulation, which results from the dependency on the gene association set rather than a single gene.

For the AJ and B6 mouse strains, we found that the number of upregulated reactions was higher than the number of downregulated reactions, whereas for the PWD mouse strain the inverse was observed (Fig. S2 lower panel). For AJ and PWD, the gene regulation followed the same trend for reaction regulation, whereas for B6 the numbers of up- and down-regulated genes and reactions remained comparable (Fig. S1).

To represents as the human NS *vs* S we compared mice that have shown high nonalcoholic steatohepatitis (NS) phenotypes (AJ mice fed with the DDC-supplemented diet) to mice that have shown high steatosis (S) phenotypes (PWD mice fed with DDC supplemented diet). For the NS *vs* S in mice (see materials and methods; AJ DDC vs PWD DDC) we identified 459 and 191 up- and down-regulated reactions, respectively (Table S 2-S3).

### Supplementary Figures

**Figure S1.**
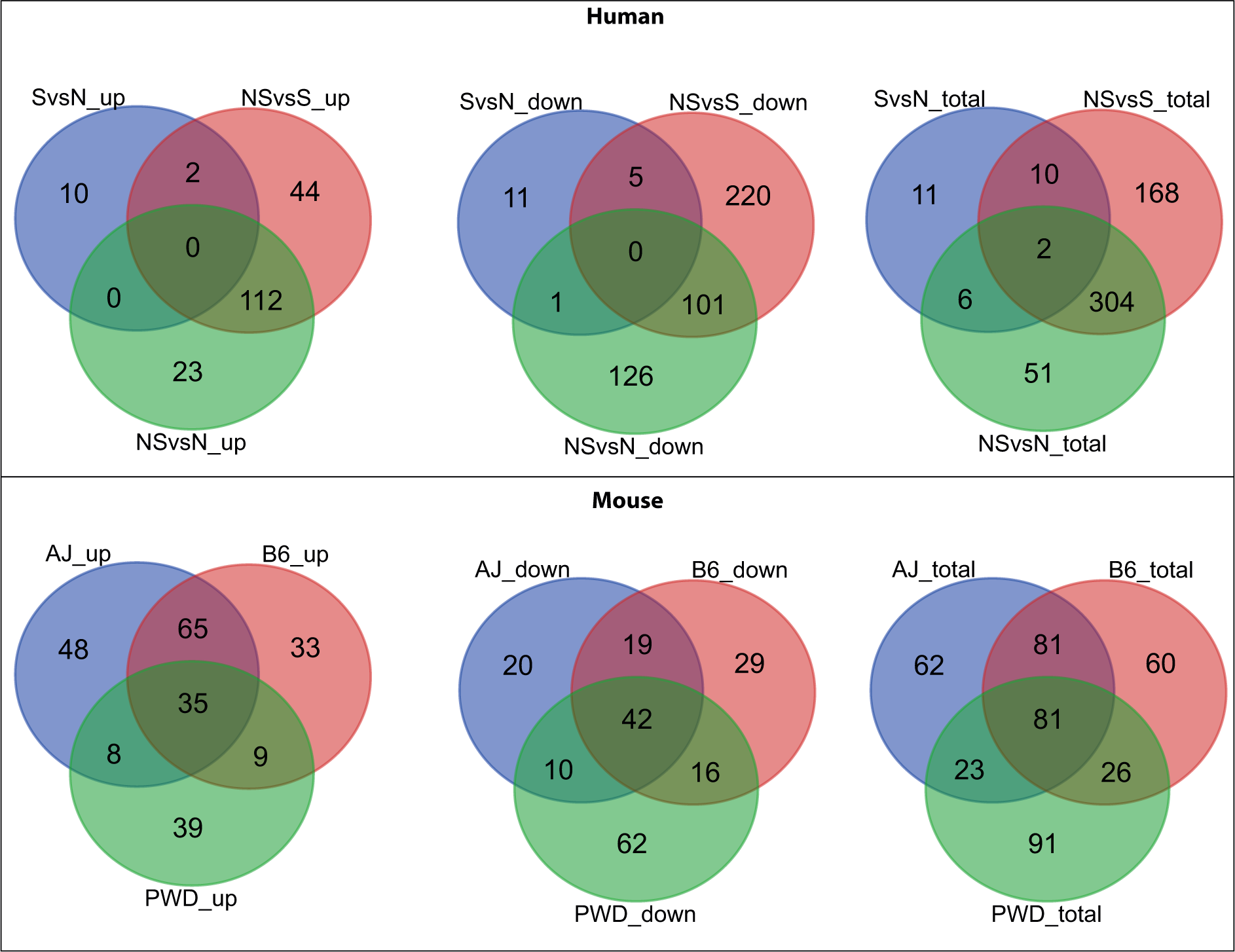
Venn diagram of differentially expressed genes of human and mouse liver samples. Upper and lower panels represent human and mouse, respectively.

**Figure S2.**
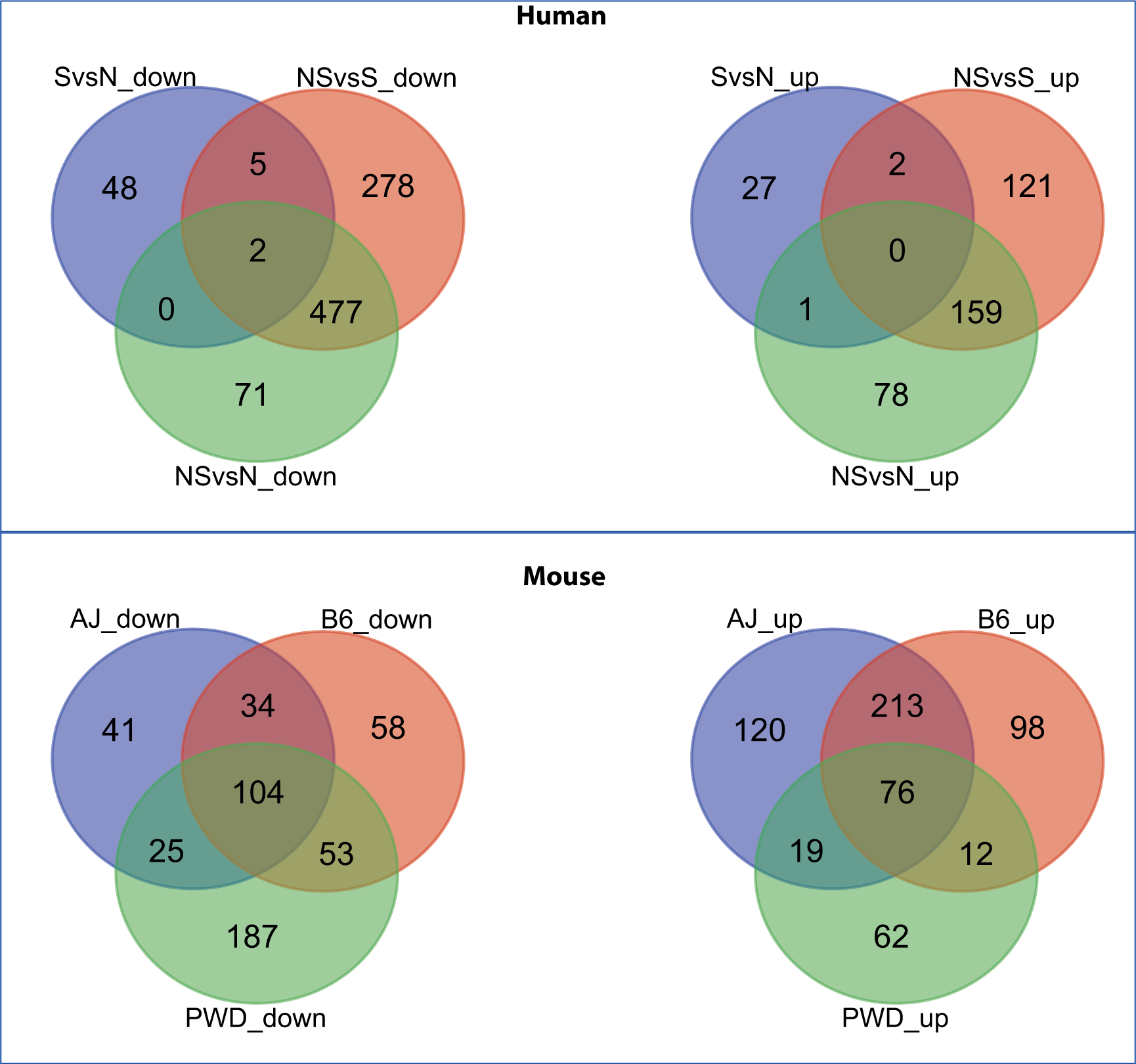
Venn diagram of up- and downregulated reactions of the iMM1415 in human and mouse liver samples. The reaction regulation metric was used to identify up- and downregulated reactions.

